# Multimodal retinal imaging by visible light optical coherence tomography and phosphorescence lifetime ophthalmoscopy in the mouse eye

**DOI:** 10.1101/2025.05.29.656845

**Authors:** Stephanie Nolen, Zhongqiang Li, Jingyu Wang, Mirna El Khatib, Sergei Vinogradov, Ji Yi

**Affiliations:** Department of Biomedical Engineering, School of Medicine, Johns Hopkins University, Baltimore, MD 21231; Department of Ophthalmology, School of Medicine, Johns Hopkins University, Baltimore, MD 21231; Department of Biochemistry and Biophysics, Perelman School of Medicine, University of Pennsylvania, Philadelphia, PA 19104; Department of Chemistry, School of Arts and Sciences, University of Pennsylvania, Philadelphia, PA 19104

## Abstract

**Significance:** Oxygen metabolism is important to retinal disease development, but current imaging methods face challenges in resolution, throughput, and depth sectioning to spatially map microvascular oxygen.

**Aim:** To develop a multimodal system capable of simultaneous phosphorescence lifetime imaging scanning laser ophthalmoscopy (PLIM-SLO) and visible light optical coherence tomography (VIS-OCT) to capture capillary-level oxygen partial pressure (pO_2_) and structural volumes in rodents.

**Approach:** C57BL/6 mice were imaged by VIS-OCT with high-definition (10 kHz raster) and Doppler (100 kHz circular) protocols. Phosphorescent probe Oxyphor 2P was retro-orbitally injected to enable intravascular PLIM-SLO imaging (200 µs pixel dwell time), while a tunable lens was used to adjust the focal depth. The extracted phosphorescence lifetimes were used for pO_2_ calculation. Simultaneous imaging utilized a shared imaging path and synchronized data collection.

**Results:** VIS-OCT images revealed detailed anatomy and Doppler shifts, while PLIM-SLO provided capillary pO_2_ at multiple depths. A hemoglobin oxygen dissociation curve related retinal arterial pO_2_ to systemic oxygen saturation as inhaled oxygen was varied. Registered simultaneous images were captured and pO_2_ was empirically adjusted for the combined excitation.

**Conclusions:** Detailed anatomical structures and capillary pO_2_ levels can be simultaneously imaged, providing a useful tool to study oxygen metabolism in rodent disease models.

## INTRODUCTION

Due to the high metabolic demands of the retina, oxygen perfusion is essential to proper visual function. Oxygen homeostasis is maintained by complex interactions between the retinal tissues and two vascular systems, the inner retinal and choroidal systems [1]. The inner retinal circulation is metabolically regulated via neurovascular coupling, similar to that in the brain [2]. The inner circulation supplies the inner retina and develops into three capillary plexuses: the superficial vascular plexus and the intermediate and deep capillary plexuses [3,4], which coincide with the retinal nerve fiber and ganglion cell layer, the inner plexiform layer, and the outer plexiform layer, respectively. The less regulated, highly vascularized choroidal circulation lies outside of the retina. Oxygen diffuses from the choriocapillaris to the outer retina through Bruch’s membrane and the tight junctions of the retinal pigment epithelium [5]. Evidence shows that there is interaction between the inner retinal and choroidal circulations, which suggests that both circulation systems warrant examination in retinal pathologies that are conventionally considered to involve either the inner or outer retina [6–8]. Pathologies may alter the microvasculature directly or change extravascular structures that impact vessels or diffusion [9]. Disruption of this delicate balance is implicated in several serious ocular diseases such as glaucoma, age-related macular degeneration (AMD), and diabetic retinopathy (DR) [5,10–13].

The significance of retinal oxygen perfusion calls for *in vivo* measurements of oxygen in the eye, yet the existing methods face outstanding challenges, particularly in localizing within tissue regions and providing depth-resolved measurements within small vessels down to the capillary level. Oxygen sensing microelectrodes provide direct ground-truth oxygen measurements of oxygen partial pressure (pO_2_) in retinal tissue and vasculature, but are invasive and only collect data at a single point [5,10,14]. Because oxygen perfusion is passive and gradient driven, the measurement of blood hemoglobin oxygen saturation (sO_2_) within the vasculature at the equilibrium state can also infer tissue oxygenation [15]. Several non-invasive imaging methods harness the oxygen-dependent hemoglobin absorption contrast to perform sO_2_ measurement.

Multi-wavelength fundus oximetry or scanning laser ophthalmoscopy are easy to adopt but are limited to 2D. These measurements are complicated by diffusive light, pigmentation, and vessel center line reflection, ultimately resulting in a lack of depth resolution. Photoacoustic ophthalmoscopy (PAOM) can image in 3D, but lacks sufficient resolution for capillaries [10], and depth resolution when applied in humans. When an ultrasound transducer is used for PAOM, physical contact with the eye is necessary and thus cumbersome.

Visible light optical coherence tomography (VIS-OCT) addresses the issue of depth resolution in previous methods, allowing spectral analysis within blood vessels to exclude confounding factors from other layers. VIS-OCT has been successfully applied in large human peripapillary vessels and validated in retinal arterioles with pulse oximetry [16]. Yet, measurements in smaller vessels around the macular region in humans and in capillaries in general are still challenging and may require additional calibrations due to less signal for averaging along the narrow radius of the vessels to smooth out the spectrum [17–19]. Furthermore, venous and capillary oximetry calculations still lack a local ground truth or gold standard for validation, since systemic pulse oximetry is only accurate for arterial blood.

The phosphorescence quenching method has been implemented to reliably measure pO_2_ in animal models, using dyes that change their phosphorescence lifetimes as a result of quenching by molecular oxygen [20,21]. Several platinum and palladium porphyrin-based dendritic oxygen probes have been designed and used in the past for *in vivo* oxygen sensing [22–24]. *In vivo* phosphorescence lifetime imaging using these probes can provide quantitative maps of oxygen partial pressure (pO_2_), excluding many confounding factors that intensity-based measures face. Wide-field imaging of the retina has been performed with such probes, but is limited primarily to large vessels due to low resolution and lack of depth control [25–28]. Two-photon phosphorescence lifetime microscopy [29] has been used to capture high resolution oxygenation points at varying depths in the eye and brain but has limited throughput to build comprehensive maps of the retina [30,31].

To address these limitations, we designed a dual-channel VIS-OCT and phosphorescence lifetime scanning laser ophthalmoscope (PLIM-SLO) rodent system to capture high-resolution volumetric tissue information and corresponding capillary-level oxygenation over a range of depths. PLIM-SLO uses point imaging and an oxygen probe, Oxyphor 2P (Ox2P) [32], to establish high-resolution spatial mapping of pO_2_ down to the capillary level, as a ground truth for VIS-OCT label-free intravascular oximetry. VIS-OCT generates high-resolution, three-dimensional structural images of the retinal tissues and can collect functional information such as blood flow through Doppler OCT. Simultaneous capabilities allow capture of spatially and temporally registered structural and vascular information. This system can be used to examine functional oxygenation and tissue structural changes in rodent disease models. The system described in this contribution leverages *in-vivo*, nondestructive imaging capability, which should be especially useful in longitudinal studies of etiology or drug effects on the retina by allowing repeated imaging of the same subject over time, without the need to sacrifice the animal for imaging. Here we demonstrate and validate the dual-channel system in mouse retinae.

## MATERIALS AND METHODS

### System design

The probe Oxyphor 2P [32] was chosen for its exceptional brightness and its ability to provide maps of absolute pO_2_ values in biological environments. Excitation light for the PLIM SLO channel was provided by an OBIS 640 nm LX diode (Coherent), to trigger high 1-photon absorption by Ox2P (peak ∼630 nm) [32] while avoiding the wavelength range of the VIS-OCT spectrometer. The excitation light was spatially filtered by a fiber, then passed through a telescope to be resized onto a tunable lens (EL-16-4-TC-VIS-20D, Optotune) located in the shared excitation/emission path. The tunable lens allowed depth scanning by changing the convergence of the collimated beam entering the mouse eye, thus changing the focal plane at the retina. It was positioned conjugate to the pupil plane and oriented vertically to avoid aberrations caused by gravity unevenly distorting the lens surface. The ∼758 nm peak emission light was separated from the excitation via a dichroic mirror with a center wavelength of 659 nm (ZT647rdc -UF1, Chroma). Phosphorescence was collected by a photomultiplier tube (PMT) (H10721-20, Hamamatsu) and converted to voltage by a transimpedance amplifier (TIA60, Thorlabs) before digitization (ATS9416, AlazarTech). A 50 µm fiber (M50L02S-B, Thorlabs) served as a pinhole in the collection path to remove the out-of-focus signal for depth sectioning. The fiber core radius was about nine times larger than the airy radius to balance the depth selection stringency with the signal strength.

**Fig. 1.**
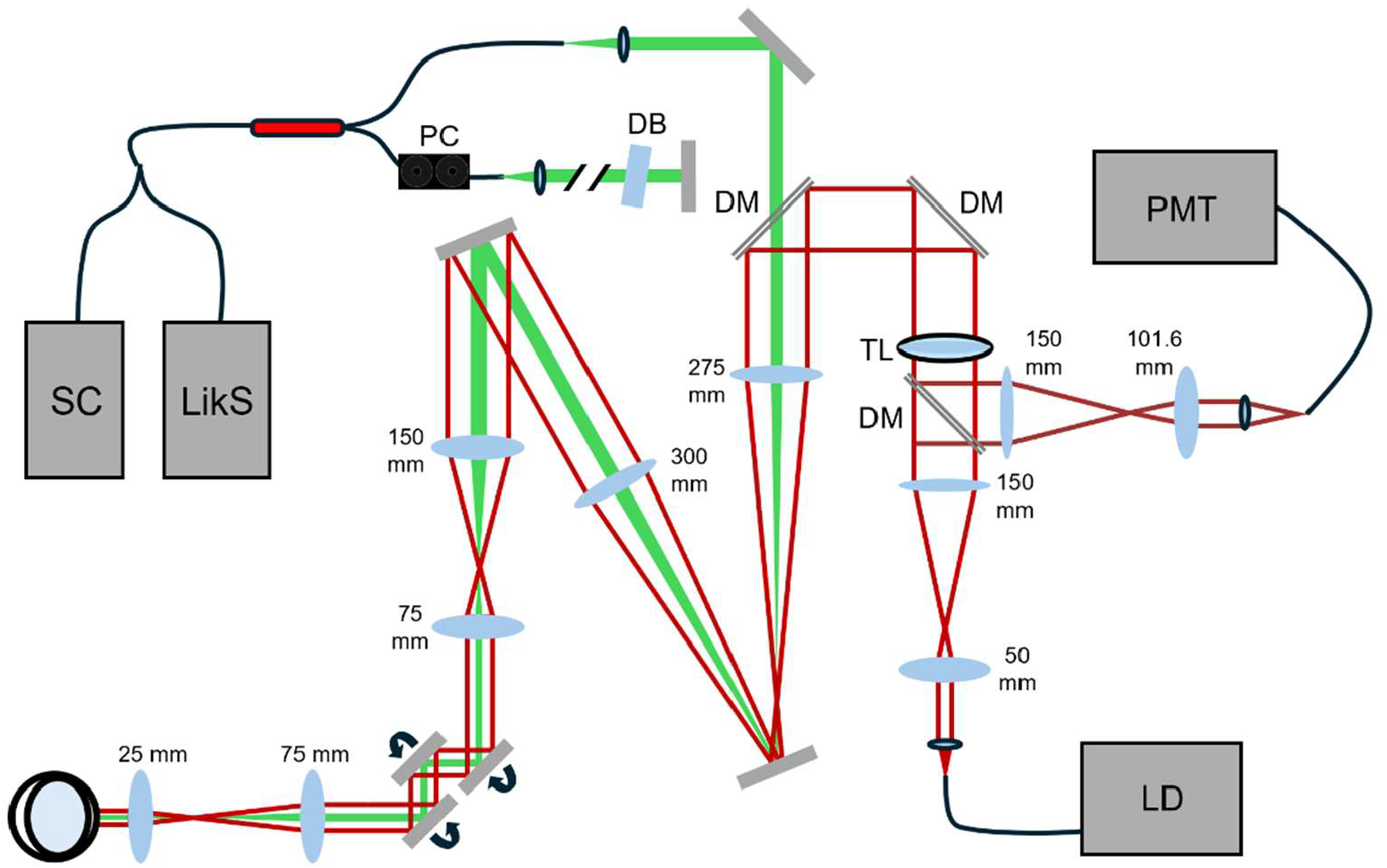
System diagram of multimodal imaging system. The VIS-OCT and PLIM are combined into a common path via dichroic mirror. An additional path is reserved via dichroic mirror for future channel development. DM – Dichroic mirror; PC – Polarization controller; DB – Dispersive blocks; SC – Supercontinuum source; LikS – Linear-in-k spectrometer; TL – Tunable lens; PMT – Photomultiplier tube; LD – Laser diode.

A SuperK Extreme supercontinuum light source (NKT Photonics) was used in the VIS-OCT channel, and a 90:10 fiber coupler (Thorlabs) was used for interferometry. The spectrum was collected by a custom visible light linear-in-k spectrometer described in Ref. [33] with a center wavelength of 559.87 nm and wavelength range of 500.07 to 635.36 nm. By collecting the spectrum linearly by wavenumber instead of wavelength, the spectrometer reduced high pass filtering of the signal by the line camera (Octoplus, Teledyne e2v) and therefore mitigated roll-off. A volume phase holographic diffraction grating (WP-1800/532-50.8, Wasatch Photonics) and F2 glass equilateral dispersive prism (PS852, Thorlabs) dispersed the light by wavenumber before it was focused onto the camera by a 75 mm objective (11-321, Edmund Optics). Small adjustments to the VIS-OCT focal plane on the retina were made by axial translation of the sample arm collimator to maximize signal. The reference arm contained glass blocks (LSM03DC-VIS and WG11050-A, Thorlabs) for coarse dispersion compensation and a paddle polarization controller (FPC020, Thorlabs) was placed on the reference fiber.

The two channels were combined in the sample arm by a shortpass dichroic mirror (T640spxr-UF2, Chroma) with a transmission band of 387-633 nm and a reflection band of 642-833 nm. The beams shared a common path and scanning mirrors (GVS001, Thorlabs; ScannerMax Saturn 9, Edmund Optics) into the mouse eye to allow physical registration of the images.

### Image Collection

After brief induction with isoflurane, male C57BL/6 mice were anesthetized by intraperitoneal injection of a ketamine/xylazine cocktail (83 mg ketamine per kg of body weight and 8.3 mg xylazine per kg of body weight). Eyes were dilated with 1% tropicamide (Alcon Laboratories, Inc.) and a custom mouse contact lens (Advanced Vision Technologies) was placed on the cornea over a thin layer of artificial tear gel (GenTeal Tears, Alcon Laboratories, Inc.) to maintain corneal moisture. The mouse was immobilized and positioned in a custom animal holder with a nosecone (Kent Scientific) modified with a bite bar for head fixation. Body temperature was maintained during imaging using a temperature controller (TC300B, Thorlabs) and flexible heating pads (TLK-H, Thorlabs) incorporated into the animal holder, with a target value of 36.7° C. Core temperature was collected via a rectal probe and thermocouple meter (20250-91, Digi-Sense) and systemic sO_2_ was collected from the hind paw using a commercial pulse oximeter (Model 8500AV, Nonin).

For PLIM-SLO imaging, mice were injected retro-orbitally in the non-imaging eye side with 0.1 mL of 200 µM Ox2P. A 256 by 256 raster scanning protocol was used, where a 10 µs pulse was used for excitation in each point, and the decay was captured for a total pixel dwell time of 200 µs. The average power incident on the cornea was 0.25 mW. The first 10 µs of signal, during the stimulation pulse, were averaged to generate phosphorescence intensity images, which were used to segment out blood vessels with the “vessel analysis” Fiji plugin [34,35]. For average vessel pO_2_ calculations, the traces from a segmented vessel were averaged, while for point-by-point maps, a 3 by 3 moving average filter was applied to the whole image first to reduce noise before calculation by averaging each trace with its neighboring traces. A low-noise oxygen map calculation was performed by taking three collections in a row and averaging the full ‘volumes’ to reduce noise at each point before performing the point-by-point processing steps.

Single exponential fitting of the phosphorescent decay signal then generated the rate constant, k, which is the inverse of the lifetime, τ. High oxygen partial pressure leads to more rapid quenching of the phosphorescence, and thus a shorter lifetime. Oxygen partial pressure was then calculated from the fitted lifetime value at each point using an independently measured Stern-Volmer-like calibration plot (see Results) that describes the relationship between the quenching of the phosphor and varying levels of oxygen partial pressure at different temperatures [28]. Mouse body temperature was maintained close to the target temperature of 36.7°.

VIS-OCT imaging was performed with 0.3 mW power at the cornea. Various protocol patterns and scanning rates up to 120 kHz can be implemented in the VIS-OCT channel. Dispersion not accounted for by the glass blocks was compensated using a time-frequency analysis based method to generate the phase correction [36]. The interferometric signal was background subtracted and Fourier transformed into volumetric images. High resolution structural images (512 × 512) of the retinal layers were collected by a slower 10 kHz line rate to reduce noise. Motion artifacts occurring during longer imaging times were compensated by a simple flattening algorithm applied to the surface of the retina. A stack of 100 flattened B-scans was then averaged to bring out fine layer details in the retinal tissue. Functional Doppler imaging was performed by scanning a dense circular pattern about the optic nerve head at a 100 kHz line rate (8192 × 32) and then subtracting adjacent A-lines to obtain the phase shift signal.

Simultaneous imaging required coordinated triggering between the two channels. Simultaneous imaging was performed with a combined imaging protocol that maintained the 200 μs pixel dwell time to allow the full phosphorescent decay period in the raster scan (256 × 256). A 50 kHz trigger signal was simultaneously sent to the line scan camera to collect 10 OCT A-lines during each SLO collection point. An additional averaging step to combine the multiple A-lines into one was included in simultaneous VIS-OCT processing. A visible light rate correction factor to compensate for VIS-OCT stimulation of Ox2P was empirically calculated and incorporated into simultaneous pO_2_ processing, as described in the Results section.

## RESULTS

### PLIM-SLO

The PLIM-SLO channel generated high resolution maps of the retinal vasculature as seen in Fig. 2. Three acquisitions were taken in rapid succession, then the phosphorescent signal during the pulse stimulation period was averaged to produce a low-noise intensity image. Fig. 2a shows an example of an image in a region adjacent to the optic nerve head, with focusing towards the surface of the retina. The phosphorescence decay traces in Fig. 2 b, taken from a cross section of points through Fig. 2 a, demonstrated a more rapid decay in the arterioles (upper and lower major vessels) compared to the venule (central major vessel). This relationship is clearly visualized in the Fig. 2 b inset, where the averaged traces of each of the major vessels showed a faster drop-off in the arterioles versus the venules. Fitting of a single-exponential function was performed on the decay traces to calculate the decay rate constant, k_Ox2P_, from the measured phosphorescent intensity I over time t:

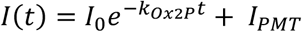

**Fig. 2.**
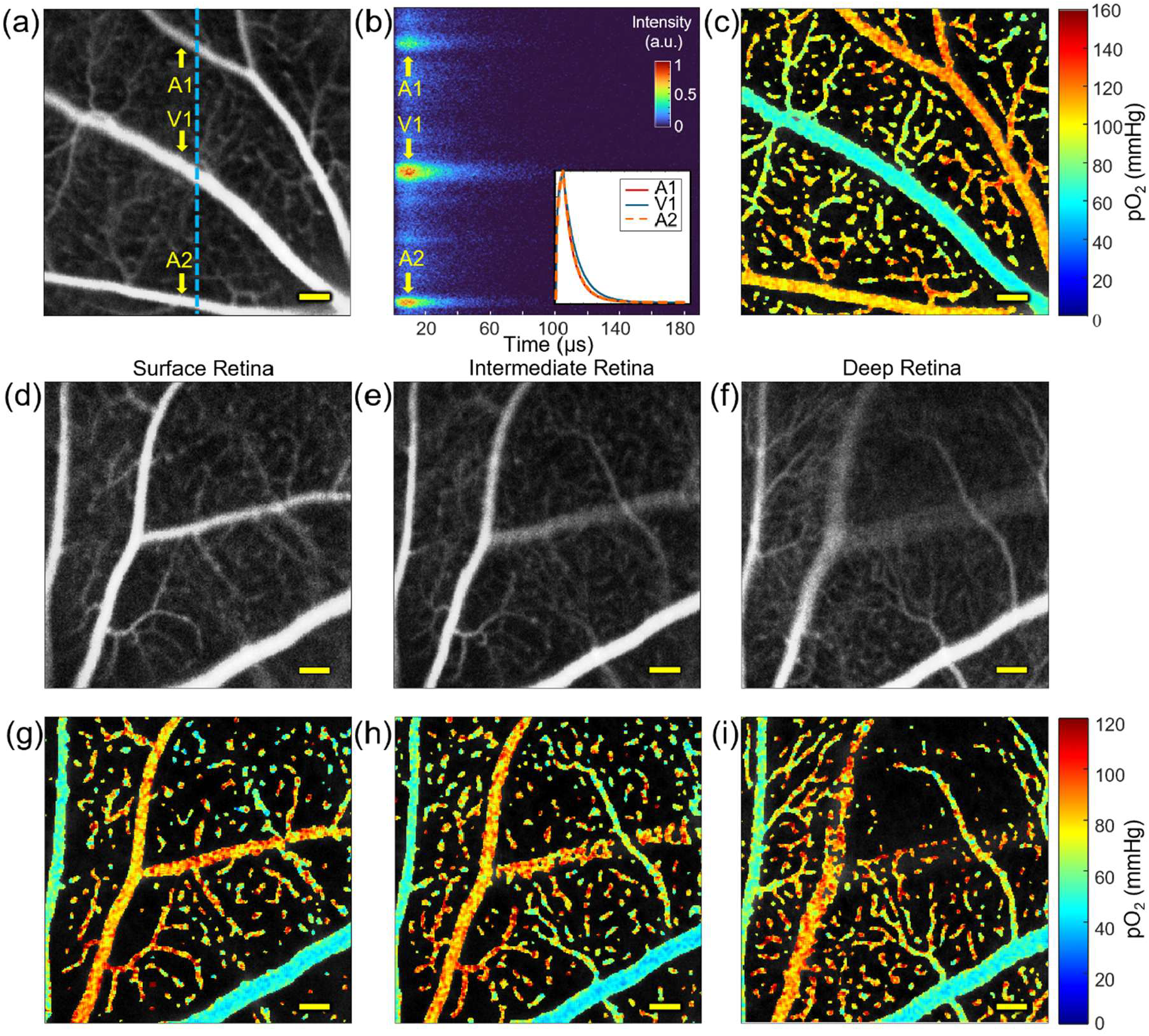
PLIM-SLO measures and maps pO_2_ *in vivo* in 3D. (a) Average of three 256 × 256 phosphorescence intensity images collected in rapid succession for low-noise imaging. (b) Normalized phosphorescence intensity traces over time from the line scan along the blue dashed line in the intensity image. Average traces from the two major arterioles and venules are displayed in the inset. (c) Point-by-point oxygen tension map of the averaged phosphorescence image with a systemic oxygen saturation of 96%. (d)-(f) 256 × 256 microvascular sections of retinal plexuses at increasing depths from the retinal surface. (g)-(i) Corresponding pO_2_ values from single collections from a mouse with systemic oxygen saturation of 89%. *All scale bars 100 μm*.

where I_0_ was the initial intensity at the start of phosphorescent decay, when t = 0, and I_PMT_ accounted for the nonzero baseline generated by PMT collection. The rate constant was then used to find the lifetime τ from the relationship τ = 1/ k_Ox2P_. The lifetime and measured temperature T were used to calculate the oxygen partial pressure pO_2_ (mmHg) using an empirical formula, as described previously [32]:

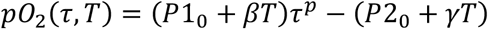

where p = -1.265777 (unitless), P1_0_ = 3.06907×10^−4^ mmHg s^-p^, β = -4.6205×10^−6^ mmHg s^-p^ °C^-1^, P2_0_ = 114.4061 mmHg, and *γ* = -1.60147 mmHg °C^-1^. After averaging the three collections in Fig. 2 a, a 3×3 moving average filter was applied to the lateral dimensions and a point-by-point calculation of the spatial pO_2_ distribution was mapped using the above equations (Fig. 2 c). Structural variations in the microvasculature over depth were demonstrated by adjusting the tunable lens through different layers of the retina (Fig. 2 d – f). Vessels located towards the retinal surface, such as arteriole branches that were in focus in Fig. 2 d, then fell out of focus in Fig. 2 f in favor of deeper structures such as venule branches. This structural organization reflected anatomy previously described in the literature of precapillary arterioles in the superficial vascular plexus and capillaries and post-capillary venules in the deep capillary plexus joined by transverse vessels and the less defined intermediate capillary plexus [37–39]. The pO_2_s of several larger vessels and branches, calculated from single collections, are shown in Fig. 2 g – i.

### VIS-OCT

High-definition VIS-OCT cross-sectional structural images of the retinal layers were generated by collecting data at a slowed 10 kHz line rate and averaging 100 flattened slices of the mouse retina along a narrow region of the slow axis (Fig. 3 a-c). In the outer retina, several bands were distinguished, including the putative inner segment/outer segment junction [40], the photoreceptor outer segment tips, the retinal pigmented epithelium, and Bruch’s membrane. An example of the functional capabilities of the device is shown in Fig. 3 d-f, where Doppler phase shifts were observed in major vessels about the optic nerve head by scanning in a 100 kHz circular pattern.

**Fig. 3.**
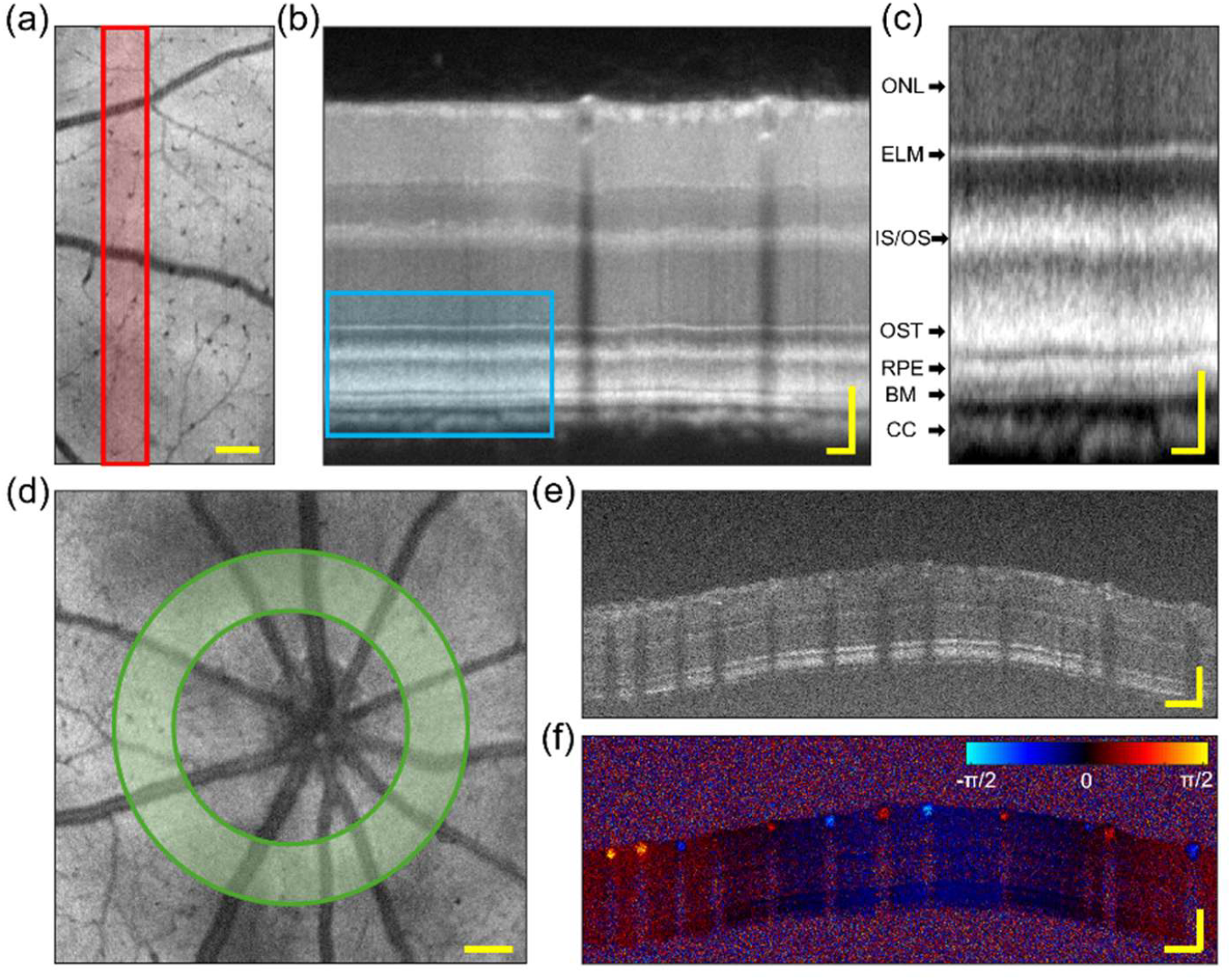
High-definition structural imaging and Doppler imaging by VIS-OCT. (a) 512 × 512 en face of the mouse retina collected at a 10 kHz line rate. *Scale: 100 μm*. (b) Flattened and averaged retinal cross section of 100 B-scans across the slow axis corresponding to the highlighted region in red. *Scales: 50 μm lateral, axial*. (c) Zoomed image of region delineated by the blue box for visualization of outer retinal layers. ONL, outer nuclear layer; ELM, external limiting membrane; IS/OS, putative inner segment outer segment junction; OST, photoreceptor outer segment tips; RPE, retinal pigment epithelium; BM, Bruch’s membrane; CC, choroid. *Scales: 50 μm lateral, 20 μm axial*. (d) 512 × 512, 100 kHz en face image of retina demonstrating circular 8192 × 32, 100 kHz Doppler scanning pattern. *Scale: 100 μm*. (e) Structural and (f) functional Doppler phase shift B-scans. *Scales: 100 μm lateral, axial*.

### Simultaneous Imaging

Simultaneous imaging was performed with a 200 μs pixel dwell time to allow the full phosphorescence decay time in the PLIM channel (Fig. 4). 10 repeats of 50 kHz VIS-OCT scans were performed at each scanned point during PLIM measurement and then averaged into one B-scan to generate an *en face* image of the retina with detailed microvasculature mapping (Fig. 4 a) that matched the simultaneously collected phosphorescence intensity image (Fig. 4 c). Retinal layer information from a stack of 50 averaged B-scans (Fig. 4 b) and spatial oxygen partial pressure distribution (Fig. 4 d) were gathered simultaneously. An empirically calculated correction factor (validated in Fig. 5 i) was applied to the fitted rate constant to cancel the effect of VIS-OCT light stimulation on pO_2_ calculation.

**Fig. 4.**
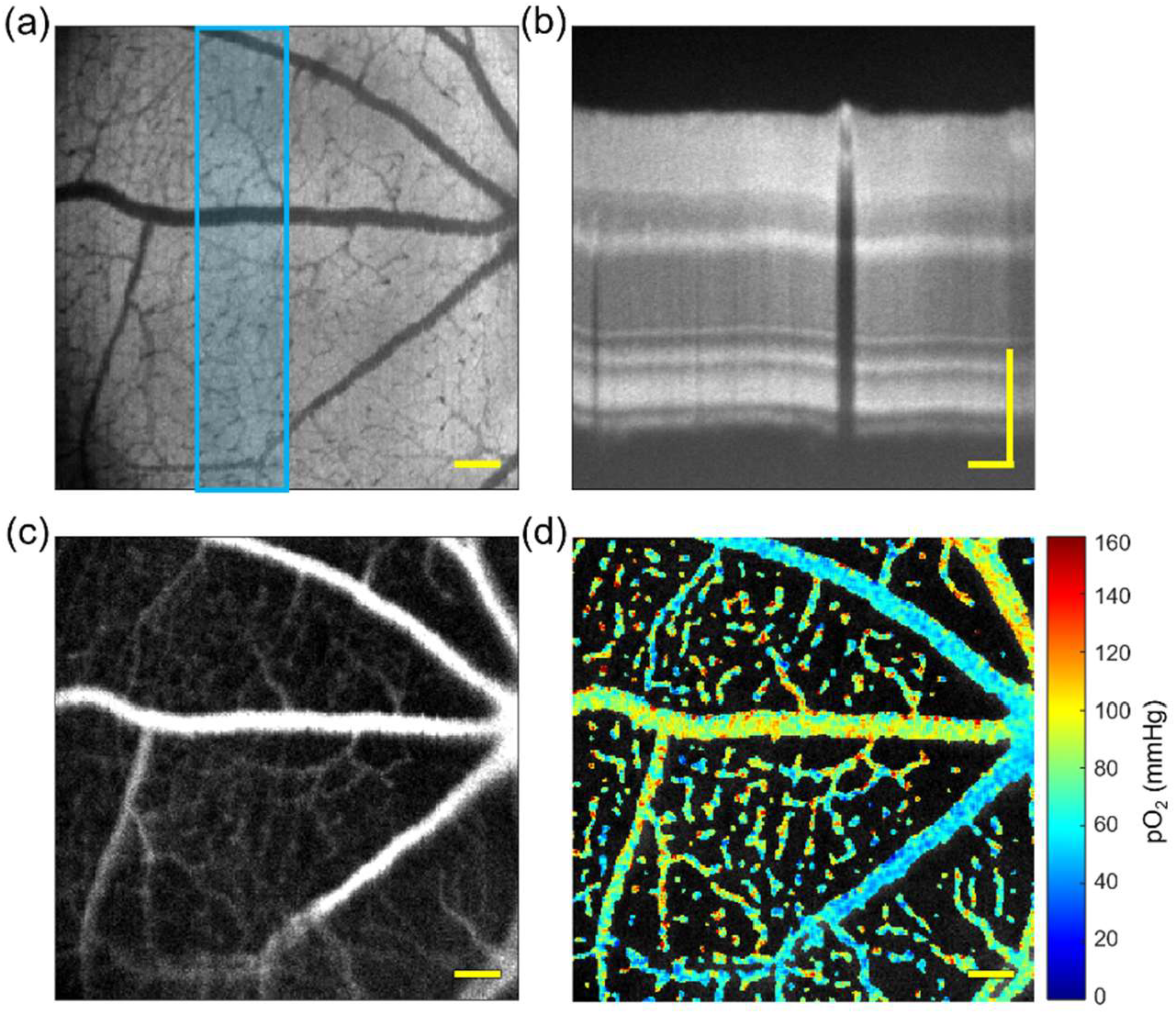
Simultaneous VIS-OCT and PLIM collection. (a) En face VIS-OCT image generated from outer portion of retina volume for increased microvasculature contrast. (b) B-scan generated from stack of 50 averaged B-scans outlined in blue on the en face. (c) Phosphorescence intensity image and (d) oxygen partial pressure map with systemic arterial saturation of 87%. *All scale bars: 100 μm*.

**Fig. 5.**
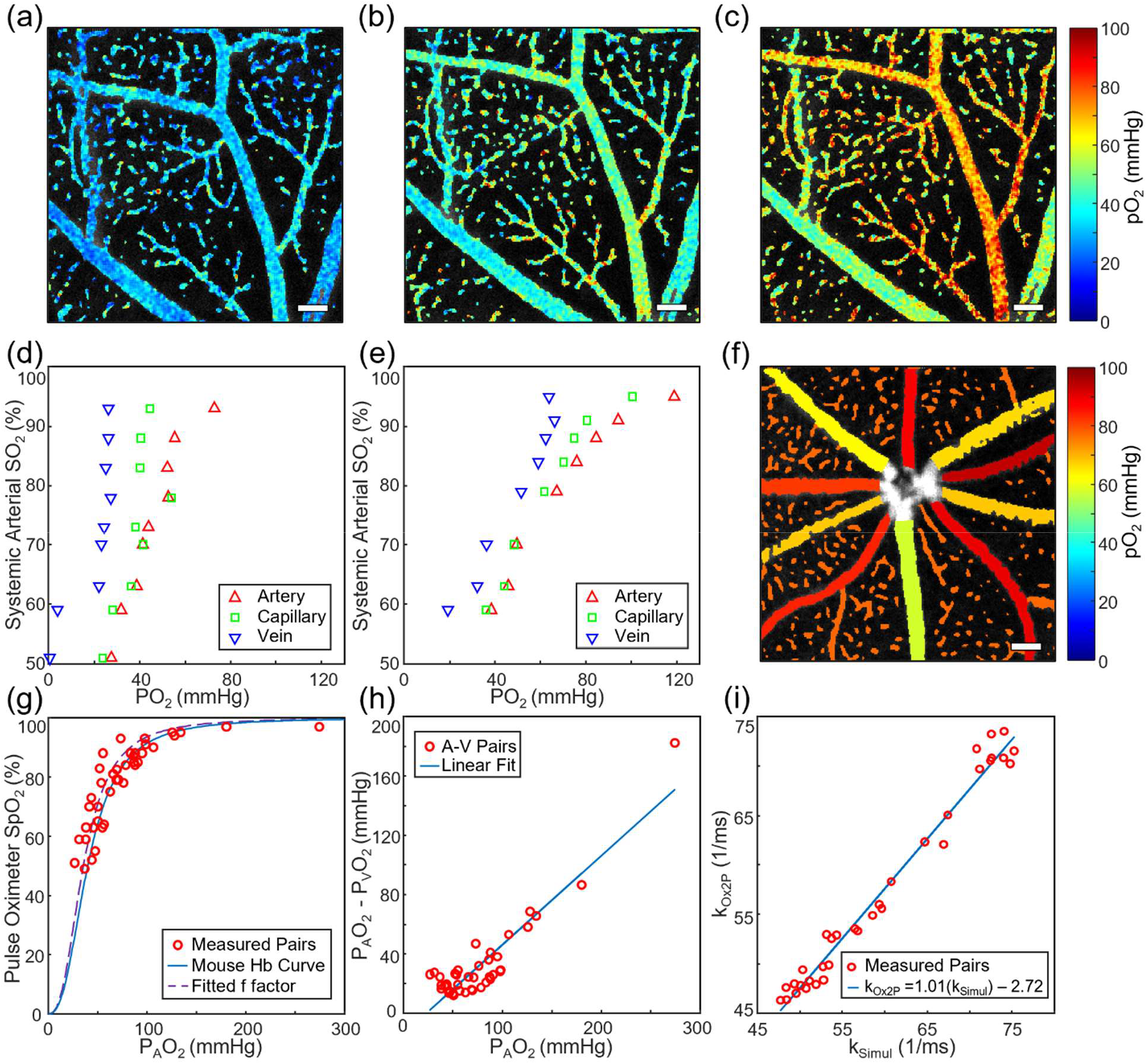
Validation of oxygen partial pressure imaging. (a)-(c) Microvasculature oxygen partial pressure maps at systemic sO_2_ values of 69%, 78%, and 88%, respectively. (d)-(e) The average oxygen partial pressure in arterioles, venules, and capillaries at several oxygen saturation states from the right eyes of two example mice in a FOV about the ONH exemplified in (f). (g) The systemic oxygen saturation and average arterial oxygen partial pressure pairs plotted with the oxygen hemoglobin dissociation curve for n = 6 mice at standard conditions and with a fitted f factor. (h) The difference between average arterial and venous oxygen partial pressure showed an increase as the average arterial partial pressure increased. (i) Fitted pairs of k values with and without OCT illumination (n = 34 vessel pairs) to calculate correction factor k_corr_ = 2.72 × 10^3^ s^-1^. *All scale bars: 100 µm*.

The visible light wavelengths supplied by the OCT channel are capable of off-peak excitation of Ox2P [32] and had to be accounted for when extracting lifetime values from the decay traces during simultaneous imaging. The measured intensity of the phosphorescence as a function of time *I*_*Simul*_*(t)* during simultaneous imaging (derived in Supplement 1) is described as:

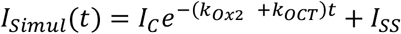

where *t* is time, where *t*=0 is the moment when the excitation pulse ends. *k*_*Ox2P*_ is the native triplet decay rate constant, and *k*_*OCT*_ is a excitation rate constant due to the background continuous wave (CW) illumination by the visible light source. The constant *I*_*C*_ is the intensity of the phosphorescence signal excited by the pulse at the start of the decay, while the constant *I*_*SS*_ is the intensity of phosphorescence due to the CW VIS-OCT illumination.

When the simultaneous images were analyzed, fitting of the phosphorescent decay generated a combined rate constant k_Simul_ equal to the sum of k_Ox2P_ and k_VIS_. To determine the native decay rate k_Ox2P_ for oxygen calculation, a correction factor k_corr_ value, equal to k_VIS_, was subtracted from the value k_Simul_, determined by fitting. k_corr_ was determined empirically by collecting sets of PLIM images with and without CW VIS-OCT illumination. It is important to mention that the molar extinction coefficient of Ox2P even between its main absorption peaks (460 nm and 630 nm) is still relatively high (∼10^3^ M^-1^cm^-1^). Therefore, given the tight spatial focusing, the excitation flux at the beam waist even at a seemingly low power of ∼0.3 mW was sufficient to increase the apparent decay rate constant by 10-20%.

### In vivo pO_2_ validation

Validation of PLIM oxygen calculations was performed by varying inhaled oxygen levels and measuring systemic sO_2_ using a pulse oximeter. The fraction of oxygen in the inhaled gas mixture was controlled via an oxygen/nitrogen gas mixer (7300 Series, MATHESON). The mixture was delivered through a snug-fitting nosecone to prevent gas leakage. Supplied oxygen was adjusted incrementally, then images were collected once systemic sO_2_ values were stable, according to the pulse oximeter for at least 15 sec without fluctuation. Changes in systemic oxygenation were reflected in the retinal pO_2_ values down to the capillary level. As visualized in Fig. 5 a-c, the 256 × 256 point-by-point calculated oxygen partial pressures increased throughout arterioles, venules, and capillaries as the proportion of inhaled oxygen and systemic sO_2_ increased.

Oxygenation calculations were quantitatively validated by collecting averaged trace values from major arterioles, venules, and capillaries within a constant field of view (FOV) as the oxygen saturation varied. In Fig. 5 d-e, sample oxygen challenges from the right eyes of two mice are shown with the average arterial, capillary, and venous values for each systemic oxygenation condition. A FOV about the optic nerve head (ONH), exemplified in Fig. 5f, was chosen to include all of the major vessels and various capillaries. Average arterial pO_2_ was higher than venous in all cases, and at most oxygen saturation levels the capillary values were located between the major vessel values (Fig. 5 d-e). At a few points, the capillary average was observed very near to or slightly higher than the arterioles.

The average arterial values during the oxygen challenge were used to generate an oxygen dissociation curve, as they were assumed to be closely related to the systemic arterial sO_2_. Six mice were imaged with a FOV about the ONH to include all major vessels for averaging. The average of the arteriole pO_2_s was paired with the systemic sO_2_ values to generate a hemoglobin dissociation curve (Fig. 5 g). To account for variations in the hemoglobin binding relationship due to temperature (T), the pO_2_s of the measured points were multiplied by a correction factor f. The correction factor, based on the Kelman equations for oxygen dissociation [41], is defined by:

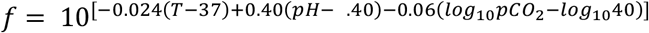

This correction factor shifts the pO_2_ of the measured points to what the pO_2_ would be for the “standard” environmental values (pH = 7.4, T = 37° C, pCO_2_ = 40) used to generate the dissociation curve. The measured temperatures were used, and a pH value of 7.4 and pCO_2_ value of 40 mmHg were assumed, as these values could not be noninvasively measured in the retina during imaging. The generated hemoglobin dissociation curve was compared to a Hill equation model from the literature, shown as a solid curve in Fig. 5g, with the C57BL/6 mouse hemoglobin Hill coefficient n = 2.59 and p50 = 40.2 mmHg [42].

The contributions from each environmental factor to *f* can easily be separated into three terms. Since the temperature-only term, f_T_, is calculated from the measured temperature, the two unknowns can be grouped into a common factor, f_pH,pCO2_, that the pO_2_ is multiplied by after temperature correction by f_T_. A least-squares fit on the mouse hemoglobin Hill equation for f_pH,pCO2_ was performed on the systemic sO_2_ and temperature-corrected pO_2_ pairs for comparison to the assumed standard pH and pCO_2_ values. The resulting fitted curve was plotted as the dashed curve in Fig. 5 g. The fitted value of f_pH,pCO2_ = 1.135 was similar to the value of f_pH,pCO2_ =1 that is generated by the standard pH and pCO_2_ values. The fitted curve for the unknown values appeared close to the literature-derived curve that assumed standard values.

The pO_2_ relationship between vessel types was further explored. The differences between the average retinal arterial and venous pO_2_s for the six mice imaged about the ONH were also calculated. The venous pO_2_s used the same combined correction factor as the arterioles from the corresponding collection. The difference between arterial pO_2_s and their corresponding venous pO_2_s showed a positive trend as the oxygen partial pressure increased (Fig. 5h), with an R-squared value of 0.84. The difference between arterioles and venules increased with hyperoxia, potentially due to metabolic regulation of the inner retina.

To calculate and validate the simultaneous imaging k correction factor, PLIM image pairs were collected with and without the 0.3 mw OCT source stimulation in rapid succession. The rate constant values from major vessels (n = 34) with (k_Simul_) and without (k_Ox2P_) stimulation were calculated from single exponential fitting of the averaged traces. The relationship was fitted by the least-squares method, with the zero-order term used to empirically define the k_corr_ value and an anticipated first order term of unity. Least-squares fitting of the pairs resulted in an R-squared value of 0.94 (Fig. 5i). The slope of the line approached unity (1.01), as expected from the theoretical relationship, and the correction factor k_corr_ was calculated to be 2.72 × 10^3^ s^-1^. The correction was applied to simultaneously collected PLIM images to provide accurate pO_2_ measurements.

## DISCUSSION

The developed instrument allowed collection of detailed structural and functional information from the rodent retina using PLIM-SLO and VIS-OCT. Spatial mapping of oxygen partial pressure at a focal plane between the superficial and deep plexuses revealed higher pO_2_ in the two major retinal arterioles and branching arterioles and lower pO_2_ in the major venule and returning draining venules. The tunable lens and pinhole were used to pass through different depths to focus on specific plexuses, with the maintenance of similar pO_2_ values in the recurring major vessels demonstrating a consistency in pO_2_ readings over time. However, there is still some overlap from background vessels visible in each plane. Depth sectioning could be improved by using a collecting fiber with a smaller diameter (acting as a smaller pinhole with a diameter closer to the airy disc of the beam as it enters the fiber) at the expense of signal loss. The fiber used was chosen to optimize phosphorescent signal strength and optical sectioning.

VIS-OCT imaging revealed high-definition structures along the retinal depth, with detailed microvasculature visible in the en face images. Diving vessels, the connections between plexuses, appeared more distinctly as dark points in this channel compared to the phosphorescence channel due to an accumulation of absorption in the axial direction through the vessel’s path perpendicular to the retinal surface. High-definition B-scans reveal detailed structural information, particularly in the outer bands of the retina. With the implementation of a circular scanning protocol, Doppler phase shift signal was collected. Doppler OCT can be used to calculate blood flow in future studies quantifying oxygen metabolism in the retina by combining flow values with the PLIM-SLO pO_2_ values.

Simultaneous imaging demonstrated physical registration of the structural VIS-OCT and phosphorescence images. The adjustment factor applied to the rate constant of the curve fitting accounted for the stimulation of Oxyphor 2P by the visible light channel. The empirically calculated relationship between the images with and without OCT stimulation had a slope close to unity, indicating good agreement with the theoretical expectation that a correction factor could be added to the fitted k for simultaneous imaging with constant background illumination. It should be noted that changing the visible light illumination (i.e. power or spectral shape) would require recalibration of k_corr_, since the constant illumination value would differ.

Physiological factors may play a role in the conversion between measured pO_2_ and oxygen saturation in the hemoglobin dissociation curve. The standard pH value of 7.4 and pCO_2_ value of 40 mmHg were assumed for the scaling factor f since noninvasive local collection in the retina was not possible during imaging. The correction factor itself was also calibrated to human hemoglobin values [41], which may lead to a slight difference in mice, but should be reasonably accurate since the Bohr effect in humans and mice have been reported as similar [42–44]. The empirical fit for f_pH,pCO2_ over the hypoxic challenge series suggested that the true values were not excessively different than the assumed standard values.

While arterial values were consistently higher than venous, there were a few instances in which capillary average values were observed very close to or slightly higher than the arterial values. This similarity between capillary and arterial values is likely due to the focal plane used during imaging. The tunable lens was used to choose a plane close to the surface of the retina, where the major vessels were well focused, which is more likely to contain highly oxygenated surface vessels branching off of the arterioles. The presence of higher oxygenated capillaries towards the surface was observed when depth sectioning through the retina as shown in Fig. 4. This distribution was also observed in the literature [30]. Arterio-arterial shunts in the superficial vascular plexus have also been reported [37], and could contribute to the high capillary average seen here.

Due to the aberrating nature of the mouse eye, there is additional room for improvement in resolution and signal quality. Incorporating adaptive optics into the reserved channel space of the device can be used to improve the resolution and signal level, particularly in the SLO channel. Improvements in the signal strength can permit a smaller pinhole for improved depth sectioning of retinal layers. Additional future development of the system hardware can improve depth sectioning further still by the implementation of remote sensing and oblique scanning of the beam into the eye for intrinsically three-dimensional PLIM-SLO imaging.

Future research using the imaging system will use the simultaneous imaging capabilities to validate VIS-OCT retinal oximetry measurements. PLIM-SLO pO_2_ measurements can be used as a gold standard value and converted to oxygen saturation using the Hill equation. Pairing with VIS-OCT spectra can be used to validate retinal oximetry calculations in all vessel sizes and types to overcome previous challenges in validating non-arterial blood. The device is also promising for studying oxygen metabolism in the mouse retina, including in disease models. The VIS-OCT channel can be used to study structural changes in the retina, while the PLIM-SLO collects coordinated functional information. Changes in slower developing diseases or drug treatment courses can be studied longitudinally by collecting information from the same subject many times over the disease course without the need to sacrifice the animal. The depth sectioning ability also allows exploration of oxygenation changes that may occur in different layers of the retina as a disease develops.

## Acknowledgement

R01EY032163, R01EY034607, R01EB034272, NSF GRFP DGE2139757, Wilmer Pool Professorship Funding. Support of the grant U24EB028941 (SAV) is gratefully acknowledged.

