## Supplement 1 for "Multimodal retinal imaging by visible light optical coherence tomography and phosphorescence lifetime ophthalmoscopy in the mouse eye"

Supplement 1: Derivation of Decay Equation with Continuous Wave (CW) Background Excitation

The rate of change in the number of the excited state triplet (T_1_) species (*n**) following the excitation pulse in the presence of the background CW illumination is described as:

$$\begin{aligned} \frac{dn^{*}}{dt}= -n^{*}{\times k}_{Ox2P}+n_{0}\times k_{OCT},\#1 \end{aligned}$$

Where *t* is time, *n*_0_ is the number of the ground state (S_0_) species, k*_Ox2P_* is the rate constant for the T_1_ depopulation due to radiative (phosphorescence) and non-radiative processes (including quenching by oxygen), *k_OCT_* is the absorption rate constant due to the CW excitation by the OCT laser, *n*^*^ is the number of probe molecules in the excited state. We consider that *n* is the total number of probe molecules in the volume, and it is constant, so that:

$$\begin{aligned} n=n^{*}+n_{0},\#2 \end{aligned}$$

We ignore the stimulated emission term, since the energy of the triplet state is significantly lower than the excitation energy, and the respective transition (T_1_→S_0_) is spin-forbidden. We also consider the transition from the initially populated excited state (S_1_) to T_1_ (intersystem crossing) instantaneous relative to the rate constants *k_Ox2P_* and *k_OCT_*.

Substituting Eqn. 2 into Eqn. 1, gives a nonhomogeneous linear differential equation:

$$\begin{aligned} \frac{dn^{*}}{dt}+\left( k_{Ox2P}+k_{OCT} \right)n^{*}=nk_{OCT},\#3 \end{aligned}$$

The complementary equation is separable, so the complementary solution n_c_^*^(t) is:

$$\frac{dn^{*}}{dt}+\left( k_{Ox2P}+k_{OCT} \right)n^{*}=0,$$

$$\int\frac{1}{n^{*}}dn^{*}= \int-\left( k_{Ox2P}+k_{OCT} \right)dt,$$

$$\begin{aligned} n_{c}^{*}\left( t \right)= Ae^{-\left( k_{Ox2P}+k_{OCT} \right)t},\#4 \end{aligned}$$

For the particular solution, the constant *n_p_^*^(t)* = B is used as the guess for the solution:

$$\begin{aligned} 0+\left( k_{Ox2P}+k_{OCT} \right)B=nk_{OCT}, \end{aligned}$$

$$\begin{aligned} n_{p}^{*}\left( t \right)=B=\frac{nk_{OCT}}{\left( k_{Ox2P}+k_{OCT} \right)},\#5 \end{aligned}$$

Therefore, the solution is:

$$\begin{aligned} n^{*}\left( t \right)= n_{c}^{*}\left( t \right)+ n_{p}^{*}\left( t \right)= Ae^{-\left( k_{Ox2P}+k_{OCT} \right)t}+\frac{nk_{OCT}}{\left( k_{Ox2P}+k_{OCT} \right)},\#6 \end{aligned}$$

Using the initial value at *t*=0, *n^*^(0)* = *n^*^_t=0_* to solve for A gives the final formula:

$$\begin{aligned} n^{*}\left( t \right)=\left( n_{t=0}^{*}- \frac{nk_{OCT}}{\left( k_{Ox2P}+k_{OCT} \right)} \right)e^{-\left( k_{Ox2P}+k_{OCT} \right)t}+\frac{nk_{OCT}}{\left( k_{Ox2P}+k_{OCT} \right)},\#7 \end{aligned}$$

Therefore, when performing simultaneous imaging, the rate constant that is calculated by the fitting, *k_Simul_*, is equal to the sum of *k_Ox2P_* and *k_OCT_*, and the phosphorescence decay is superimposed on a constant background:

$$\begin{aligned} I_{Simul}\left( t \right){=I}_{C}e^{-\left( k_{Simul} \right)t}+I_{SS},\#8 \end{aligned}$$

$$\begin{aligned} k_{Simul}= k_{Ox2P}+k_{OCT}, \#9 \end{aligned}$$

The constant *I_C_* is the intensity of the phosphorescence signal excited by the pulse at the start of the decay, while the constant *I_SS_* is the intensity of phosphorescence due to the CW VIS-OCT illumination.

The excitation rate constant *k_OCT_* can be estimated by considering:

$$\begin{aligned} k_{OCT}=\sigma\times\Phi,\#10 \end{aligned}$$

where *σ* is the molecular excitation cross-section at the wavelength of the OCT laser and *Φ* is the photon flux through the excitation volume. The molecular excitation cross-section is proportional to the molar extinction coefficient *ε*, which for Ox2P is on the order of 10^2^-10^3^ M^-1^cm^-1^ near 500 nm. The cross-section *σ* is found as:

$$\begin{aligned} \sigma= \frac{\ln\left( 10 \right)\times1000\times\varepsilon}{N_{a}},\#11 \end{aligned}$$

where *N_a_* is the Avogadro number. The photon flux is defined as:

$$\begin{aligned} \Phi= \frac{P\times\lambda}{h\times c\times S},\#12 \end{aligned}$$

where *P* is the OCT laser power at the focus (i.e. attenuated due to tissue absorption and scattering), *λ* is the excitation wavelength, *h* is the Planck constant, *c* is the speed of light in the medium, and *S* is the beam cross-section at the waist.

While the exact value for *P* in the focal volume is difficult to measure/estimate, attenuation of the power, compared to the measured incident power under the objective, by 5-10 times, due to the absorption/scattering, results in the excitation rate constant values constituting 10-20% of the native decay constant *k_Ox2P_* at physiological oxygen pressure of 30 mmHg. These estimates are in good agreement with experimentally measured correction values *k_corr_* (see main text).
